# The antimicrobial peptide cathelicidin is critical for the development of Th17 responses in severe inflammatory disease

**DOI:** 10.1101/2022.01.27.477976

**Authors:** Katie J Smith, Danielle Minns, Brian J McHugh, Rebecca K. Holloway, Richard O’Connor, Anna Williams, Lauren Melrose, Rhoanne McPherson, Veronique E. Miron, Donald J Davidson, Emily Gwyer Findlay

## Abstract

Multiple Sclerosis (MS) is a highly prevalent demyelinating autoimmune condition; the mechanisms regulating its severity and progression are unclear. The IL-17-producing Th17 subset of T cells has been widely implicated in MS and in the mouse model, experimental autoimmune encephalomyelitis (EAE). However, the differentiation and regulation of Th17 cells during EAE remain incompletely understood. Although evidence is mounting that the antimicrobial peptide cathelicidin profoundly affects early T cell differentiation, no studies have looked at its role in longer term T cell responses. Now, we report that cathelicidin drives severe EAE disease. It is released from neutrophils, microglia and endothelial cells throughout disease; its interaction with T cells potentiates Th17 differentiation in lymph nodes and Th17 to exTh17 plasticity and IFN-γ production in the spinal cord. As a consequence, mice lacking cathelicidin are protected from severe EAE. In addition, we show that cathelicidin is produced by the same cell types in the active brain lesions in human MS disease. We propose that cathelicidin exposure results in highly activated, cytokine-producing T cells which drive autoimmunity; this is a mechanism through which neutrophils amplify inflammation in the central nervous system.

## Introduction

In this project we examined the differentiation of cytokine-producing T cells during Multiple Sclerosis (MS). MS is a demyelinating, neurodegenerative disease of the central nervous system (CNS) (1). Through the modelling of MS using experimental autoimmune encephalomyelitis (EAE), we now understand that T cells play a central role in driving this disease (2-8). However, the priming of pathogenic T cells during MS is complex and is still not fully understood. In particular, the longer-term behaviour of T cells in the CNS, and how other innate and adaptive immune cells present can influence their behaviour, is unclear.

Like in other autoinflammatory conditions (9), the Th17 subset of cells is particularly important for driving disease in MS and EAE (10-14), through their ability to cross into the CNS following priming in the lymph nodes (15-18); once there, Th17 cells also drive disease through contributing to blood brain barrier (BBB) breakdown by attracting MMP-releasing neutrophils to the site (19-22).

Neutrophils have a dynamic relationship with T cells and it is well understood that they can influence T cell activation and migration (23, 24). There is now substantial evidence that neutrophils are important and pathogenic in EAE and MS. Neutrophil populations expand during both diseases and move into the CNS (25-27). The peripheral neutrophil populations are also dysregulated, with an activated phenotype (28). Depletion of neutrophils abrogates EAE disease (4, 29, 30). The precise mechanisms through which neutrophils worsen EAE and increase severity of autoimmune conditions have not, however, been described; and while Th17 cell impact on neutrophils in the CNS is known (4) the reverse - the impact of neutrophils on Th17 cell differentiation and survival - is still unclear.

We have previously shown that neutrophil release of the antimicrobial host defence peptide cathelicidin, which occurs during degranulation and release of extracellular traps, potentiates Th17 differentiation *in vitro* and in models of acute inflammation (31). However, its role in longer-term inflammation or in inflammation of the CNS is not known.

We now demonstrate for the first time that cathelicidin is not expressed in the healthy central nervous system or secondary lymphoid organs but is strongly produced during EAE and in the active demyelinated brain lesions of patients with MS. Cathelicidin production plays a fundamental role in disease pathogenesis. Mice lacking cathelicidin have reduced incidence of EAE as T cells production of pro-inflammatory cytokines, and in particular Th17 cell plasticity to IFN-γ production, is reduced. We propose that cathelicidin is critical for the development of pathogenic longer-term Th17 responses in inflammatory disease.

## RESULTS

### Cathelicidin is expressed by multiple cell types in lymphoid organs and the central nervous system

Cathelicidin is released in lymph nodes during acute inflammation, yet whether this occurs during longer-term inflammation is unknown. To assess this, we used the chronic inflammatory model, experimental autoimmune encephalomyelitis (EAE), a model of autoimmune-mediated demyelination in Multiple Sclerosis. We induced disease with the myelin peptide MOG^35-55^ in wildtype (WT) C57BL/6JOlaHsD mice, following the standard protocol, and tracked signs of disease for 28 days. Mice showed consistent onset of disease (SuppFig1A) with mean day of symptom presentation being day 12 and median score being 2 (complete lack of tail mobility, altered gait and loss of balance). Inflammation and immune cell infiltration in the spinal cord increased consistently (SuppFig1B) and T cells were detected in the spinal cord from day 7 post-immunisation. The numbers of the total T cell population (SuppFig1C,D) and IL-17 producers (SuppFig1E) peaked at day 14, which is also the peak of symptoms, in agreement with previously published data (13, 32).

**Figure 1:**
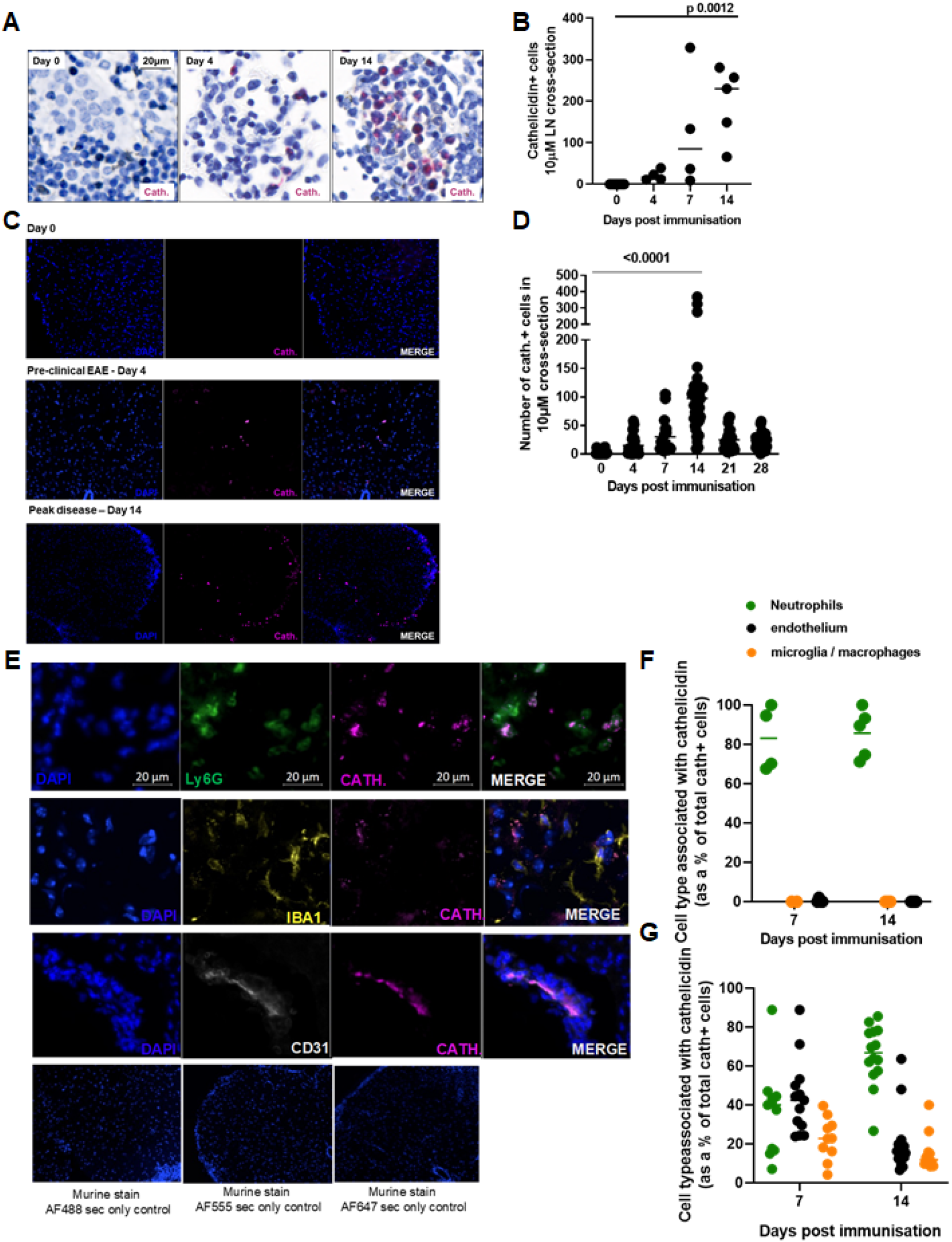
The antimicrobial peptide cathelicidin is produced in the central nervous system during EAE. EAE was induced in WT C57BL/6J mice on day 0 and mice observed for 28 days. Throughout the experiment mice were humanely culled, perfused with 4% paraformaldehyde and cathelicidin-producing cells were quantified in (A, B) draining inguinal lymph nodes and (C,D) spinal cords. (E) On day 14 co-stainng was performed to determine production of cathelicidin by Ly6G+ neutrophils, Iba 1+ microglia / macrophages, and CD31+ endothelial cells. Contribution of each cell type to overall cathelicidin production in the (F) draining inguinal lymph node and (G) spinal cord was determined on day 14 post immunisation. Data shown are individual data points with line at median. Statistical tests used: B− one way AN OVA, D − two-tailed t test comparing day 14 to day 0. N values: B- 5; D − 13-42 sections analysed from 4-6 mice; F - 5: G − 13-18 sections from 4 mice.

As we hypothesised that exposure of these T cells to cathelicidin would promote their differentiation into Th17 cells and increase their pathogenicity, the first step was to establish whether and when cathelicidin exposure occurred, as it has not previously been characterized throughout a chronic inflammatory *in vivo* model. We therefore examined the inguinal lymph node, which drains the injection site, and the spinal cord, the site of inflammation.

In steady state mice cathelicidin was not detected in lymph nodes (Fig1A,B) or spinal cord (Fig1C,D). However, as the immune response to immunisation developed, increasing amounts of cathelicidin were detected. We noted low level detection in the inguinal lymph nodes as early as day 4 post-immunisation and it increased with time, with high levels of detection on day 14 (peak of disease) (Fig1B).

In the spinal cord, cathelicidin was detected from day 4 post immunisation, consistent with previous observations that neutrophils pass into the CNS very early after immunisation (4, 25, 30, 33). While cathelicidin-positive cells appeared during the preclinical stage of the disease, they were substantially increased during the acute phase (Fig1C,D). This is, to our knowledge, the first observation of cathelicidin in the murine spinal cord.

Having established that cathelicidin is expressed in the spinal cord, we next sought to determine which cells were releasing it. Cathelicidin is known to be inducible in a variety of cell types. It is predominantly produced by neutrophils and released from the secondary granules by degranulation (34, 35); it is also present on extracellular traps (36). However, it is also expressed by other cells, including T cells (37, 38) adipocytes (39), epithelium (40-42) and mast cells (43) in mice and humans. In the inguinal lymph node, cathelicidin was produced almost entirely by neutrophils (Fig1A, F) with evidence of some having been released, as it was not associated with cells. This agrees with our previous work demonstrating that cathelicidin in the lymph nodes, following inoculation with heat-killed *Salmonella typhimurium*, is released from neutrophils (31). However, cathelicidin in the murine spinal cord was detected in a variety of cell types. The majority of cathelicidin was associated with Ly6G^+^ neutrophils (∼70% of expression was from neutrophils on day 14) (Fig1E,G), but some expression was noted in CD31^+^ endothelial cells (∼15% of expression) and Iba1^+^ microglia / macrophages (∼15% of expression).

Together, these data demonstrate that cathelicidin is widely produced during EAE, primarily by neutrophils, and that it persists longer term over the course of disease, with expression peaking at maximal disease severity.

### Cathelicidin is expressed during human Multiple Sclerosis disease

Next, we wished to determine whether cathelicidin is also expressed in human MS. To do so we analysed post-mortem brain tissue from 7 patients from the Edinburgh Brain Bank (patient information listed in supplementary table 1). Expression of cathelicidin was noted, surprisingly, in control brain tissue as well as in MS brain tissue (Fig2A,B) – the first demonstration, to our knowledge, of cathelicidin expression in the healthy human CNS. We noted that this expression was localised and next quantified cathelicidin in separate lesions, using the International Classification of Neurological Diseases guidelines. Lesions develop from normal-appearing white matter. Active lesions are characterised by an influx of immune cells, activated resident microglia and demyelination. Lesions may undergo active remyelination but can also become chronically inactive where there are few inflammatory cells and extensive demyelination. Moreover, lesions can become chronically active with a heavily demyelinated core and activated glial cells at the rim (44-46). We noted a significant increase in cathelicidin expression in the active demyelinated lesions (Fig2C). In contrast, the chronic active, chronic inactive, and re-myelinating lesions did not have increased cathelicidin compared to control tissue (Fig2C).

**Figure 2:**
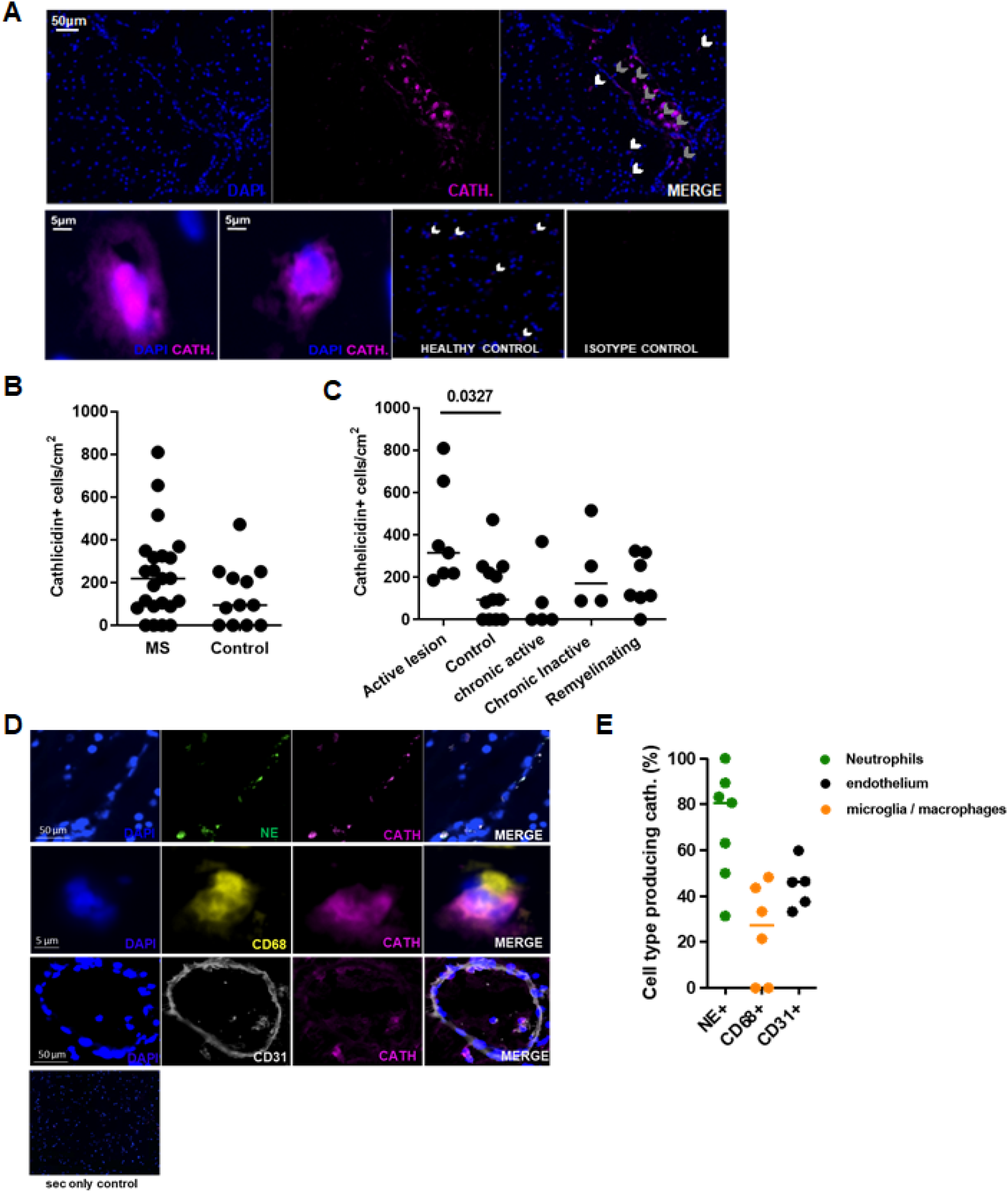
Cathelicidin is produced in the healthy and inflamed human brain. Post mortem brain sections were obtained from patients who died of Multiple Sclerosis and control patients with no neurological disease. (A-C) Cathelicidin (Cath.) production was quantified using immunofluorescent imaging. Grey arrowheads indicate cathelicidin within blood vessels and white arrowheads indicate cathelicidin outside vessels. (D) Co-staining was performed to determine production of cathelicidin by neutrophil elastase (NE)+ neutrophils, CD68+ microglia / macrophages or CD31+ endothelial cells. (E) Contribution of each cell type to overall cathelicidin production was determined in the active lesions. Data shown are (8, C, E) individual data points with line at median. Statistical tests used: (C) two tailed t test. N values: 8 and C − 7 patients in MS group and 3 patients in control group, each data point represents an area of the brain analysed; E − 3 patients, each data point represents a lesion analysed. Images in (A) and (D) are representative of at least 3 patients.

In the active lesions the same cell types co-localised with human cathelicidin as we had observed in EAE – namely neutrophils (neutrophil elastase, NE+), microglia / macrophages (CD68+), and endothelial cells (CD31+) (Fig2D, E). Quantification of active lesions was performed to identify the proportion of cathelicidin contributed by each cell type; the pattern in these lesions was roughly similar to murine samples, with the majority of the cathelicidin coming from neutrophils and a substantial minority from endothelial cells and microglia / macrophages.

Together, these data show that cathelicidin is expressed by a variety of cell types in the human and murine central nervous systems during chronic neuroinflammation. To our knowledge, this is the first example of production of any antimicrobial peptide in control human brain tissue without neurological disease.

### Cathelicidin promotes severe disease in EAE

Next we examined whether expression of cathelicidin is required for disease, by inducing EAE in mice lacking cathelicidin (*Camp*^*-/-*^, knockout, KO), first produced in (47). Observation of clinical signs of illness demonstrated that KO mice consistently showed significantly attenuated disease compared to WT mice (Fig3A,B).

**Figure 3:**
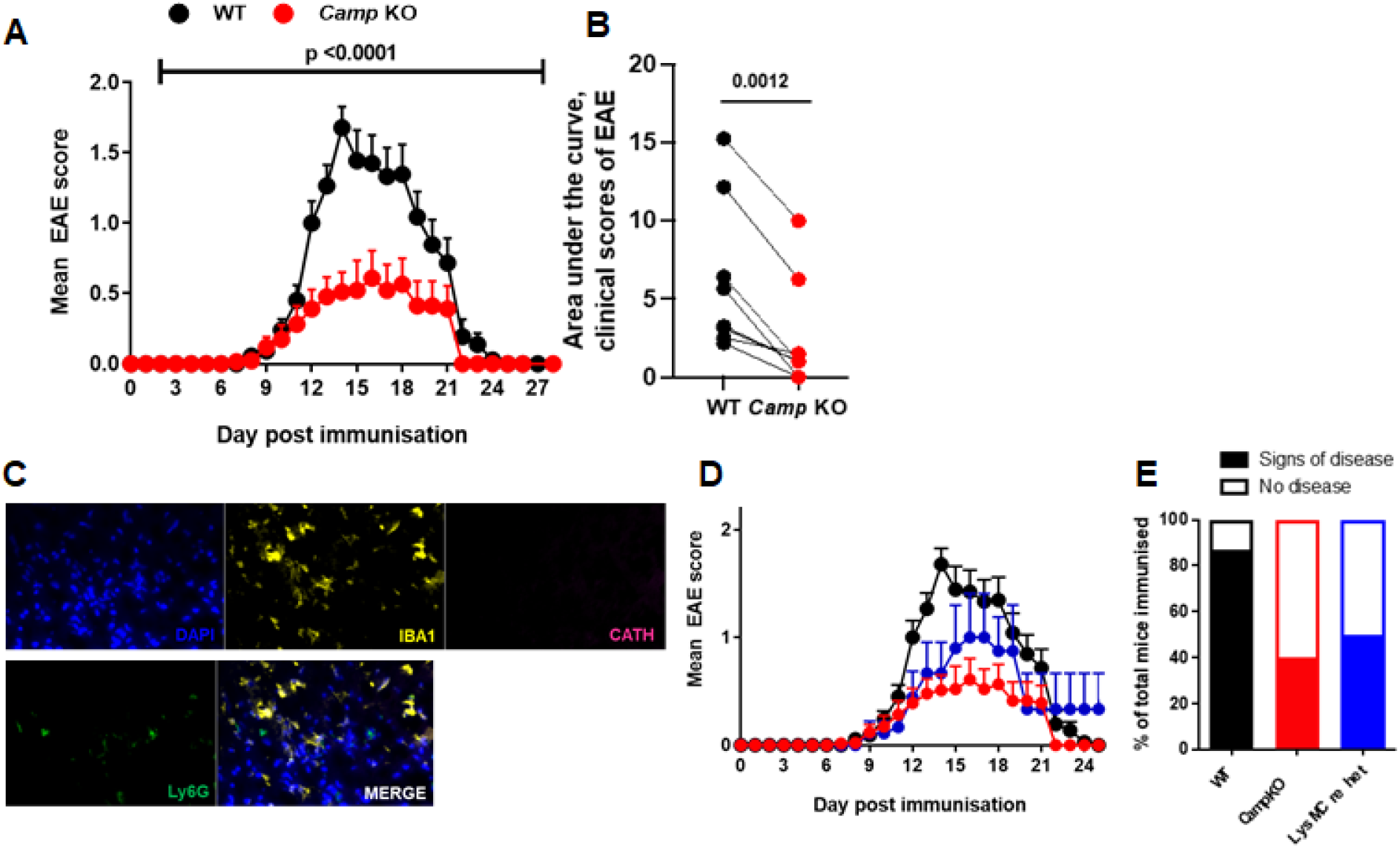
Mice lacking cathelicidin are resistant to EAE disease. EAE was induced in WT C67BL/6J and cathelicidin knockout (*Camp KO*) mice on day 0. (A) Clinical signs of illness were noted throughout the experiment and (B) area under the curve calculated for each individual experiment (C-E) Conditional knockout LysMCre mice lacking cathelicidin in myeloid-lineage cells were generated (see SuppFig2) and (C) immunofluorescence performed to confirm microglia and neutrophils lacked cathelicidin. (D,E) clinical signs of disease were tracked in conditional knockout mice. Data shown are (A.D) mean and standard error. N values: WT 85. *Camp*^*-/-*^ 71, LysMCre 8. Images in C are representative of 3 mice. Statistical tests used A −one way ANOVA, B − paired two tailed t test.

Give the multiple cell types known to be capable of expressing cathelicidin, particularly in the context of inflammation, we wanted to start to define the key cellular compartment for cathelicidin production affecting EAE disease severity, so generated a new conditional cathelicidin knockout mouse. This LysMCre mouse lacks cathelicidin in myeloid-lineage cells (generation described in SuppFig2). This mouse had no cathelicidin in microglia or neutrophils (Fig3C). Observation of clinical signs of illness over time (Fig3D) and incidence of disease (Fig3E) showed the conditional KO mice phenocopied the full KO animals and had attenuated incidence of disease, demonstrating that myeloid cell production of cathelicidin is sufficient for full disease penetrance and therefore that the noted endothelial cell production is not essential for development of disease.

### Cathelicidin does not affect cell infiltration to the spinal cord but increases T cell activation

Having shown that cathelicidin is important for EAE development, we next examined its mechanism of action. We have previously demonstrated that cathelicidin induced survival of T cell subsets (31) and others have shown it to be a chemoattractant for T cells (48). As T cells drive pathology in EAE (16, 18, 49-52), we therefore hypothesised that fewer T cells would be present in the spinal cord of KO mice, and that this was the cause of the reduced disease severity. To test this, we carried out a full flow cytometric phenotyping of spinal cord immune cells over time in both WT and KO mice. To our surprise, there were no observable differences in leukocyte infiltration between WT and KO mice during the EAE timecourse (Fig4A,B). In addition, the relative proportions of different subsets of immune cells detected in the spinal cord were not significantly different between the genotypes (Fig4C,D). To further confirm these findings, we performed immunofluorescent staining for CD3^+^ T cells in WT and KO spinal cord and found no differences in number or location on day 14 post-immunisation (Fig4E,F).

**Figure 4:**
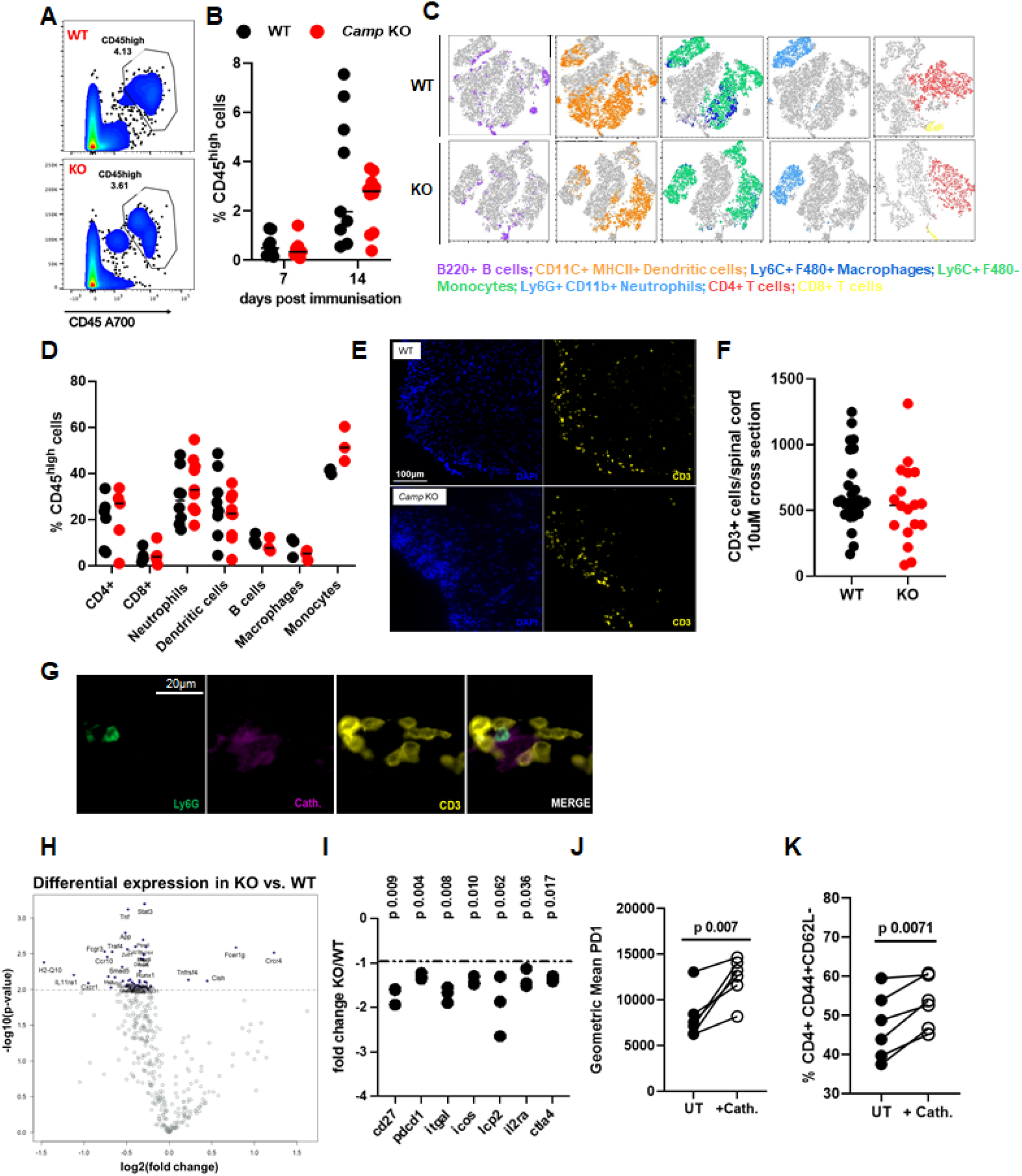
Mice lacking cathelicidin are resistant to induction of EAE despite normal infiltration of spinal cord T cells. EAE was induced in wildtype (WT) C57BL/6J and cathelicidin knockout *(Camp*^*-/-*^, KO) mice on day 0. (A, B) on days 7 and 14 CD45^high^ cell infiltration into the spinal cord was quantified by flow cytometry. (C, D) Immune cell subsets in the CD45^high^ gate were delineated on day 14 in the spinal cord then (E,F) CD3+ T cell numbers in the spinal cord were confirmed on day 14 by immunofluorescence. (G) Neutrophil release of cathelicidin was observed in close contact with T cells in the spinal cord. (H,I) On day 7 draining inguinal lymph nodes were removed and CD4+ T cells isolated. Gene expression differences between WT and KO T cells were assessed on a Nanostring mouse immunology chip (J,K) Splenic CD4+ T cells from naïve WT mice were incubated with CD3/CD28 stimulation and 2.5μM synthetic cathelicidin for 24 hours before activation was assessed by flow cytometry. Data shown are individual data points with (8, D, F) line at median. Statistical tests used: (I) two-tailed t tests with multiple comparison correction; (J,K) paired two tailed t test. N values: A − 8-10; B - 8-10; D − 3-8 F − 18-29 sections from 3 mice; I − 3, J. 6 K-6. Images are representative of 3 mice.

Together, these data show that cathelicidin is widely expressed during EAE and plays an important, non-redundant role in disease pathogenesis. However, this was not secondary to a decrease in the total number of T cells or other immune cells entering the CNS.

As the overall magnitude of immune cell flux into the CNS was similar in WT and KO mice, and spinal cord T cell numbers at the peak of disease were the same, we therefore hypothesised that the phenotype of the T cells, and the cytokines they produced, were altered in the absence of cathelicidin. Production of pro-inflammatory cytokines from T cells is of critical importance to the development of severe EAE disease (18, 53-55). We have previously shown that cathelicidin strongly preferentially potentiates Th17 but not Th1 or Th2 differentiation during acute responses (31) and so we hypothesised the same phenomenon was occurring during EAE, particularly as we observed that T cells in the lymph nodes and spinal cord (Fig4G) were exposed to cathelicidin throughout disease.

To examine this, we decided to characterise T cells in the draining lymph node, just before they move into the spinal cord and initiate observable disease. We isolated CD4^+^ T cells from the inguinal node of WT and KO mice on day 7 post-immunisation, and analysed gene expression differences between the strains of mice, using a Nanostring Immunology chip. A large number of genes were altered between WT and KO T cells (Fig4H).

Firstly, T cells isolated from the inguinal lymph nodes of KO mice (therefore differentiating in the absence of cathelicidin after induction of EAE) were less activated than those from WT mice. Expression of genes encoding CD27, PD1, LFA-1, ICOS, and LCP2 were significantly lower in KO CD4^+^ T cells (Fig4I). To confirm these findings at the protein level and to examine if this was a direct effect of cell exposure to cathelicidin, we incubated isolated splenic CD4^+^ T cells with synthetic cathelicidin *ex vivo*, alongside CD3/ CD28 stimulation. In this experiment we observed, agreeing with the Nanostring data, a significant up-regulation of PD1 (Fig4J) and CD44 expression, and a loss of CD62L (Fig4K) following exposure to cathelicidin. These data suggest that cathelicidin is able to increase T cell activation.

### Cathelicidin drives Th17 cell differentiation and plasticity

Next, we noted that many genes encoding proteins which are part of the Th17 differentiation pathway were down-regulated in KO mice. These included the TGF-β receptors, STAT3, RORγt, the aryl hydrocarbon receptor AHR and SOCS3 (Fig5A). The large number of genes modulated by the absence of cathelicidin suggest this is a key pathway through which cathelicidin mediates its functional impact, supporting our previous work in this area. Importantly, showing specificity in the immunomodulation, genes relating to the Th1 and Th2 pathways were not altered in KO mice (Fig5B), demonstrating *in vivo* that cathelicidin does not suppress all cytokine production indiscriminately but specifically affects Th17-related genes.

**Figure 5:**
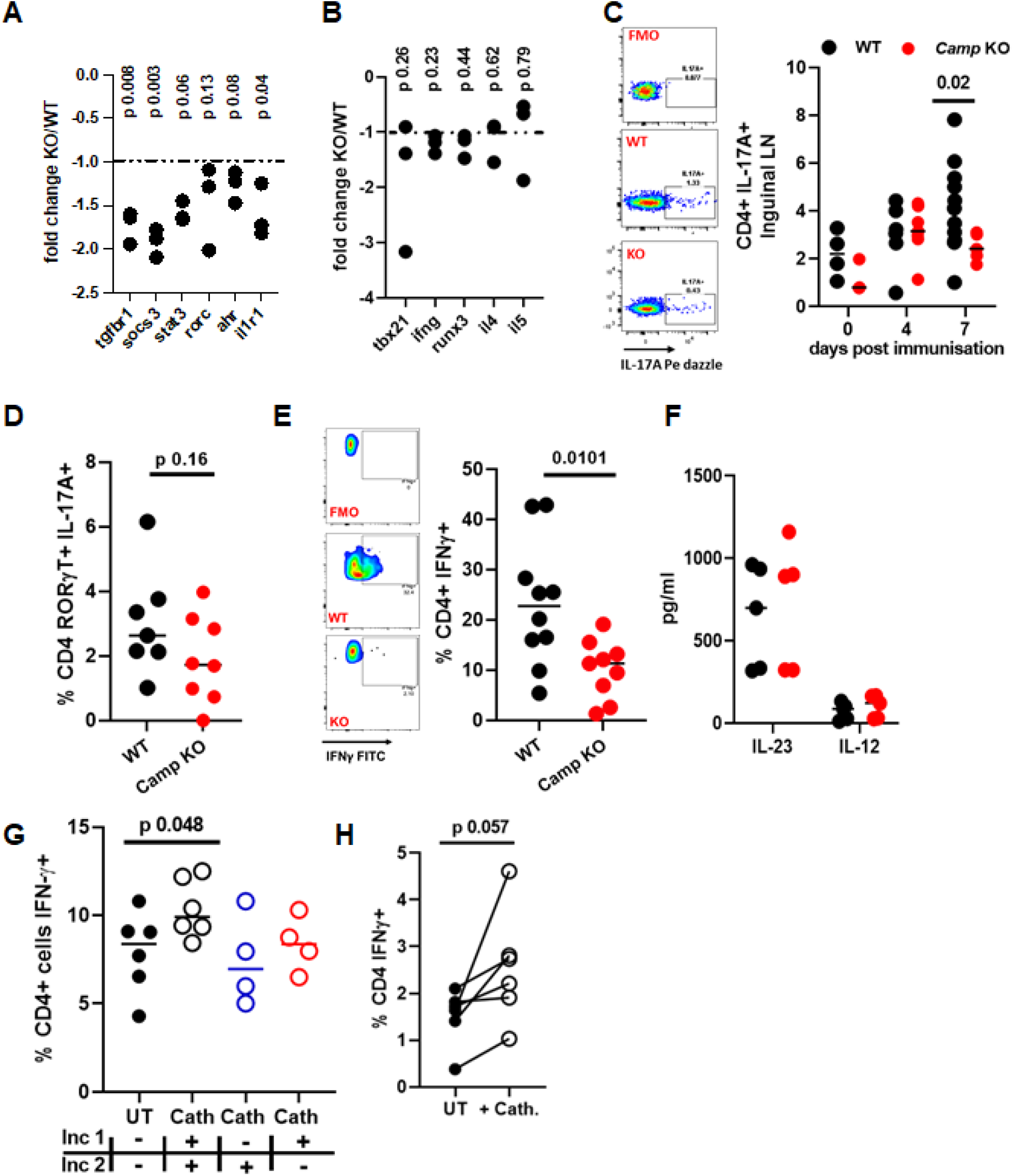
In the absence of cathelicidin T cell differentiation into Th17 cells and to exTh17 cells is impaired. EAE was induced in WT C57BL/6J and cathelicidin knockout *(Camp*^*-/-*^, KO) mice on day 0. On day 7 draining inguinal lymph nodes were removed and CD4+ T cells isolated. (A-B) Gene expression differences between WT and KOT cells were assessed on a Nanostring mouse immunology chip and (C) On day 7 IL- 17 production was assessed by flow cytometry in the inguinal lymph nodes. (D,E) on day 14 cytokine production was assessed by flow cytometry or (F) by ELISA in the spinal cord. (G) WT splenic CD4+ T cells were cultured with Th17 driving medium for 48 hours (incubation 1, inc 1) then recombinant IL-12 for 72 hours (incubation 2, inc 2) before cytokine production was assessed by flow cytometry. (H) 0 n day 7 of EAE inguinal lymph nodes were removed from WT mice and incubated for 72 hours with IL-12 and synthetic cathelicidin. Data shown are individual data points with lines at median. N values: A, B -3; C- 3-11; D − 7-8; E − 9-10; F · 5; G − 5 for WT and Cath ++, 4 for Cath -+ and +-; H− 6. Statistical tests used: (A,B) − 2 tailed t tests with post-hoc correction for multiple comparisons; (D, E)· two tailed t test; (G, H) − paired two tailed t test.

To examine this at the protein level, we isolated CD4^+^ T cells from the inguinal node. The production of IL-17A was indeed significantly suppressed in mice lacking cathelicidin compared to WT animals at day 7 in the disease course (Fig5C), but not at baseline, indicating that KO mice do not demonstrate this defect prior to an immunological challenge. CD4^+^ T cells established a basal level of IL-17A production in response to EAE induction, regardless of genotype (Fig5C); however, when the Th17-biased immune response amplified at day 7 in WT mice, the same amplification did not occur in KO mice. This was associated with protection from severe pathology, in concordance with multiple previous studies demonstrating that a reduction in IL-17 production leads to less severe EAE disease (12, 13, 56).

IL-17-producing Th17 cells are, however, not the whole story in EAE pathogenesis. There is considerable plasticity of Th17 subsets depending on the cytokines to which they are exposed (11, 57-62). In particular, Th17 cells (RORγt^+^, producing IL-17) which are exposed to IL-12 or to IL-23 up-regulate IFN-γ production in a STAT4 and Tbet-dependent fashion (57, 58). Therefore, during EAE, early RORγt^+^ IL-17A^+^ cell populations in the draining lymph node lose IL-17A and become a pathogenic IFN-γ^+^ ‘ex-Th17’ population (11, 60-64) - indeed, almost all the T cell IFN-γ production in the spinal cord is from cells which were once IL-17 producers (11). Early suboptimal Th17 potentiation in the lymph node, in the absence of cathelicidin, could modulate this process, with a consequent lessening of the frequency of these cells in the CNS. We hypothesised that cathelicidin potentiates EAE pathology not only by enhancing early Th17 differentiation but also by influencing later Th17 to exTh17 conversion.

To test this, we looked at cytokine production in the spinal cord on day 14 of EAE. Th17 cell numbers in the spinal cord were reduced at peak disease, but not significantly (Fig5D). Instead, in agreement with our hypothesis, we noted a significant reduction in IFN-γ production by CNS CD4^+^ T cells (Fig5E).

The conversion of Th17 cells to IFN-γ producers occurs following IL-12 or IL-23 signals (57, 58, 63, 65). It was possible that cathelicidin KO mice had less of these cytokines in the CNS, but we confirmed that this was not the case with spinal cord wash ELISAs (Fig5F). Therefore we proposed that cathelicidin is able to potentiate the response to the available CNS cytokines in Th17 cells in the same way it potentiates responses to TGF-β in naive T cells.

To investigate this we cultured WT splenic T cells in Th17 differentiation medium for 48 hours in the presence or absence of 2.5 μM cathelicidin. Cells were then washed and re-incubated with IL-12 to promote IFN-γ production, again in the presence or absence of cathelicidin. We found (Fig5G) that cathelicidin strongly promoted IFN-γ production in cells which had previously been in Th17 driving conditions.

We wondered at which point cathelicidin signals are important for this conversion to IFN-γ producers. To answer this, two stages of cathelicidin exposure were provided-either in the initial Th17-driving conditions or in the later IL-12 medium (Fig5G). We found that if cathelicidin was included in one of these stages no boosting of IFN-γ production was seen - but if it was included in both stages, significant enhancement of IFN-γ occurred. This suggests that the full range of cathelicidin-producing cells described in Figures 1 and 2 may contribute to this phenotype, with neutrophils initially promoting Th17 differentiation in the lymph nodes and priming for transition to ex-Th17 cells, but also spinal cord neutrophils and microglia / macrophages potentially driving this pathogenic T cell plasticity at the site of damage.

Finally, we investigated whether T cells in the draining lymph nodes of mice during early EAE would enhance their IFN-γ production if they were exposed to cathelicidin *ex vivo*. We immunised WT mice and removed the draining inguinal lymph nodes on day 7 post-immunisation. We re-stimulated lymph node single cell suspensions with MOG peptide in the presence of IL-12, to induce conversion of antigen-specific T cells (Fig 5H). Exogenous cathelicidin added in to this process enhanced IFN-γ production, indicating that exposure to cathelicidin in the spinal cord would be expected to further enhance this conversion to ex-Th17 cells.

Together, these data indicate that the generation of a pathogenic ex-Th17 population of T cells in the CNS is impaired in the absence of cathelicidin signalling. These results lead us to propose a model in which cathelicidin has three impacts on the generation of pathogenic T cells in EAE. Firstly, it directly increases the activation of T cells. Next, it specifically potentiates Th17 differentiation and IL-17 production in the lymph node. Finally, it enhances Th17 to exTh17 differentiation and IFN-γ production in the spinal cord.

## DISCUSSION

Here we demonstrate that the antimicrobial peptide cathelicidin drives severe autoimmune disease in the mouse model of Multiple Sclerosis. Specifically, exposure of CD4^+^ T cells in the draining lymph node to neutrophil cathelicidin enhances their differentiation into Th17 cells. When these cells move into the central nervous system, they are further exposed to cathelicidin released by neutrophils, microglia and endothelial cells and this potentiates Th17 differentiation into IFN-γ producing ‘exTh17’ cells. This work extends previous studies showing that neutrophils are important in autoimmune disease by providing a mechanism through which they can drive inflammation – the specific potentiation of IFN-γ-producing exTh17 cells in the CNS.

Cathelicidin is widely expressed in autoimmune and autoinflammatory conditions (66) as well as during infections, and plays a key role in driving inflammation in these diseases – including chronic obstructive pulmonary disease (67), psoriasis (68, 69) and atherosclerosis (70). A number of pro-inflammatory mechanisms of action of cathelicidin have been described including activation of the NLRP3 inflammasome (71) promoting uptake of self nucleic acids (36, 66, 68), and enhancement of cytokine and chemokine release. To these we now add specific promotion of pathogenic T cell differentiation and long-term influence on adaptive T cell cytokine production.

We demonstrate here that cathelicidin is expressed in the healthy human brain, in which it was produced by neutrophils, microglia / macrophages and endothelial cells. The purpose of this cathelicidin release is currently unknown; to our knowledge, this is the first demonstration of any antimicrobial peptide being produced in the human brain in the absence of neurological disease. We also demonstrated that cathelicidin was widely released in the inflamed mouse spinal cord during EAE and active MS lesions in human brain. It has not previously been described in MS or EAE although expression has been noted in the olfactory bulb, cerebellum, medulla oblongata and spinal cord during experimental meningitis expression (72). The expression of cathelicidin in our data, during sterile inflammation, agrees with previous work showing its release in other systems can be triggered by mediators released during sterile inflammation – such as leukotriene B4 (73) or phenylbutyrate (74) - as well as strictly infection-triggered mediators. A greater knowledge of the role of antimicrobial peptides in sterile inflammation and in the CNS in particular is required as we now begin to understand the wide-ranging effects these peptides have on the immune system.

We observed co-localisation of cathelicidin with microglia and endothelial cells as well as neutrophils. However, we have not conclusively determined production in the EAE / MS system. It is difficult to rule out co-localisation owing to nearby neutrophil de-granulation releasing cathelicidin and it being taken up by other cells. We believe that microglia and endothelial cells are actively producing cathelicidin owing supporting evidence from previous studies; endothelial production of cathelicidin has been noted in meningitis infections of mice (75) and microglial production has been demonstrated *in vitro* (76-78). Consequently, the question of which cells are important for releasing cathelicidin into the vicinity of CNS T cells remains open. We have attempted to answer this question with our conditional knockout lysMcre mice. EAE in these mice – which do not have cathelicidin in their microglia or neutrophils but do in their endothelial cells (and in other cells such as mast cells and epithelium) has an almost identical disease course to full knockout mice. This implies that the neutrophil cathelicidin in the lymph node and microglial cathelicidin in the spinal cord are key but that endothelial cathelicidin is unimportant. Relative contributions from microglia and neutrophils are currently unknown and are the focus of current research.

The differentiation of Th17 cells in the draining inguinal lymph node and its enhancement by cathelicidin agrees with our previous data (31) where we demonstrated this phenomenon occurring in the 72 hours following inoculation with heat-killed pathogens and with influenza virus. Now, we extend those studies by demonstrating this neutrophil-driven Th17 potentiation occurs in longer term inflammatory models.

The exTh17 conversion experiments performed *ex vivo* demonstrate that both early (lymph node) and late (CNS) cathelicidin is important for the final differentiation of IFN-γ-producing cells. Our finding here, that cathelicidin induces IFN-γ production in cells previously incubated in Th17-driving conditions, is surprising. In our previous studies, we found that cathelicidin incubated with naïve T cells does not affect IFN-γ production (positively or negatively) in Th1-driving conditions, and it suppresses IFN-γ in cells which are actively receiving TGF-β signals (both of which were published in (31)). These disparate outcomes on cytokine production in naïve and activated T cells, in different cytokine milieu, demonstrate the complexity of these systems. Unravelling the signalling pathways involved will be critical. Interestingly, the further differentiation of spinal cord Th17 cells into exTh17 cells (11, 58) has been shown to be dependent on the aryl hydrocarbon receptor (11). Our previous studies demonstrated that initial enhancement of Th17 cells by cathelicidin is likewise AhR-dependent, and we hypothesise that a potentiation of AhR signalling in the spinal cord is occurring in T cells which come into contact with released cathelicidin there.

Overall, this study describes antimicrobial peptide expression in the central nervous system and establishes the role of cathelicidin in directing pathogenic T cell responses in long term inflammation. This work extends previous studies showing that neutrophils are important in autoimmune disease by providing a mechanism through which they can drive inflammation – the specific potentiation of IFN-γ producing cells in the CNS.

## Author contributions

Conceptualization: KJS, DJD, EGF; Funding acquisition: EGF, DJD, VM; Investigation: KJS, DM, Ro’C, BMcH, LM, RMcP, RKH, EGF; Methodology: VM, AW EGF; Project administration: EGF; Resources: AW, DJD, EGF; Supervision: EGF; Writing – original draft: KJS, EGF; Writing – review and editing: KJS, VM, AW, DJD, EGF.

## Acknowledgments

We thank the Queen’s Medical Research Institute flow cytometry team (Shonna Johnston, Will Ramsay, Mari George), SURF histology team, and the Institute of Genetics and Cancer Nanostring Team (Alison Munro) for help and advice. We thank Dr Anne Astier, Dr Robert Gray and Professor Julia Dorin for helpful discussions and Professor Steve Anderton for advice on the EAE model. We would like to thank Matt Sharp, Julie Thomson, Ailsa Travers and the University of Edinburgh Veterinary services team for help with mouse generation, breeding and maintenance.

## Methods

### Mice

Wild type (WT) C57Bl/6JOlaHsd and cathelicidin knockout (*Camp*^*tm1Rig*^, KO) mice were bred and housed in individually ventilated cages, under specific pathogen-free conditions. Female mice between 9-13 weeks of age were used for EAE experiments; for *in vitro* cell stimulations both male and female mice between 6 and 12 weeks were used. We have previously determined no sex differences exist in Th17 potentiation by cathelicidin. KO mice were backcrossed onto WT stocks for 10 generations. Mice were housed in accordance with the ASPA code of practice for the UK; this includes temperatures from 19-24°C, humidity from 45-65%, and a 12 hour light / dark cycle.

All animal experiments were performed by fully trained personnel in accordance with Home Office UK project licences PAF438439 and 70/8884, under the Animal (Scientific Procedures) Act 1986. This project licence outlined the program of work and was approved by the University of Edinburgh Animal Welfare Ethical Review Body. Each experimental protocol was approved by the University of Edinburgh veterinary team.

### EAE

WT and KO 9-to 13-week-old female C57BL/6J mice were injected on day 0 subcutaneously in both hind legs with 140ug Myelin Oligodendrocyte Protein^35-55^ (MOG) emulsified in Complete Freud’s Adjuvant (CFA), and on day 0 and 1 intra-peritoneally with 100ul of 50ng/ml pertussis toxin in PBS (Hooke Laboratories, #EK-2110). Mice were scored every two days until day 6, and then daily based on a an EAE scale: 0 – no disease, 1 – flaccid tail, 2 – impaired gait and/or impaired righting reflex, 3-substantially impaired gait and 4 – partially hind leg paralysis. Mice with a grade 2 or above were provided with hydrated food on the floor of the cage. Mice with grade 4 were culled immediately.

### Murine tissue and single-cell preparations

Mice were culled by perfusion with PBS and exsanguination by severing of major vessels. Single-cell preparations of spleens and lymph nodes were achieved by mashing the tissues through a 100μM strainer and washing with PBS. For the spleen red blood cells were lysed using RBC Lysis Buffer, as per the manufacturer’s instructions (BD Biosciences, #555899). Preparations of brain and spinal cord were achieved by transferring whole brain and spinal cord to a glass Dounce with 2ml HBBS and manually homogenising for 50 passes of the Dounce. Cell suspension was centrifuged, and the cell pellet resuspended in 1ml FBS and 33% Percoll, and 1ml 10% FBS was layered on top and spun for 15 minutes at 800xg at 4°C with no break. Cells were washed and resuspended in PBS ready for staining.

### Flow cytometry

Cells were stained for surface markers for 30 minutes at 4°C, protected from light. Intracellular cytokines were assessed by incubating cells for 4 hours at 37°C with Cell Stimulation Cocktail containing protein transport inhibitors (eBioscience, #00-4970-03). Cells were fixed, permeabilised and stained for cytokines using BD Cytofix/Cytoperm (BD Biosciences, #554722) as per the manufacturer’s guidelines. Cells were fixed, permeabilised and stained for transcription factors using the True-Nuclear Transcription Factor Buffer Set (Biolegend, #424401). Cytokines and transcription factors in the same FACS panel were stained with the FOXP3 transcription factor staining kit (eBioscience, #00-5523-00), as per the manufacturer’s instructions. Cell viability was assessed by flow cytometry fixable live/dead yellow (ThermoFisher #L34959) or Zombie NIR Fixable Viability Kit (Biolegend, #423105).

### Flow cytometry antibodies

CD45 (clone 30-F11, Biolegend, #103127, 1:200), CD45 (30-F11, BD Biosciences, #564225, 1:200) CD4 (GK1.5, Biolegend, #100406, 1:200), CD4 (GK1.5, Biolegend, #100453, 1:200), CD8 (53-6.7, Biolegend, #100741, 1:200), IL-17A (TC11-18H10.1, Biolegend, #506938, 1:100), IL-17F (9D3.1C8, Biolegend, #517004, 1:100), IFNγ (XMG1.2, Biolegend, #505826, 1:100), CD11c (NF18, Biolegend, #101226, 1:200), CD11b (M1/70, Biolegend, #101212, 1:150), Ly6G (IA8, Biolegend, #127618, 1:200), I-A/I-E (M5/114.15.2, Biolegend, #107635, 1:300), F4/80 (BM8, Biolegend, #123117, 1:200), Ly6C (HK1.4, BD Biosciences, #128021, 1:150), B220 (RA3-6B2, Biolegend, #103245, 1:150), T-bet (4B10, Biolegend, #644805, 1:100), RORγT (B2D, eBioscience, #17-6981-80, 1:100) CD121a (JAMA-147, Biolegend, #113509)

### NanoString

WT and *Camp*^*-/-*^ KO mice were immunised as previously described. On day 7 post-immunisation mice were perfused with 10ml PBS and inguinal lymph nodes were processed as above. DAPI-CD3+CD4+ T cells were sorted using a BD FACSAria™ II (BD Biosciences) and RNA was extracted immediately using the Qiagen RNAeasy Mini Kit (Qiagen, #74104), as per the manufacturer’s guidelines. Multiplex gene expression analysis (Mouse Immunology Panel) was performed by HTPU Microarray Services, University of Edinburgh. Data analysis was performed using nSolver 4.0 and nCounter Advanced Analysis software.

### Peptides

Synthetic murine cathelicidin (mCRAMP) (GLLRKGGEKIGEKLKKIGQKIKNFFQKLVPQPEQ) was synthesized by Almac (Penicuik, Scotland) using Fmoc solid phase synthesis and reverse phase HPLC purification. Peptide identity was confirmed by electrospray mass spectrometry. Purity (>95% area) was determined by RP-HPLC and net peptide content determined by amino acid analysis. Lyophilized peptides were reconstituted in endotoxin free water at 5 mg/ml. Reconstituted peptides were tested for endotoxin contamination using a Limulus Amebocyte Lysate Chromogenic Endotoxin Quantitation Kit (Thermo Scientific, UK #88282).

### *In vitro* plasticity experiments

Whole splenocytes were prepared by mashing the tissue through a 100uM strainer and washing with PBS. Red blood cells were lysed using RBC lysis buffer, as per the manufacturer’s instructions (BD Biosciences, #555899). 150,000 cells were plated per well of round-bottom 96-well plates in complete medium (RPMI, 10% foetal calf serum, 10 units/ml penicillin, 10 μg/ml streptomycin and 2 mM L-glutamine, all supplied by Gibco, ThermoFisher UK). All cells were differentiated for 48 hours in the presence of plate-bound αCD3 (2.5ug/ml; Biolegend, #100339), rIL-6 (20 ng/ml; Biolegend, #575706), rIL-23 (20 ng/ml; Biolegend, #589006) and rTGFβ (3 ng/ml; Biolegend, #580706), with or without 2.5μM mCRAMP. Media was carefully removed and replaced with either rIL-12 (25ng/ml; Biolegend #575402), with or without 2.5μM synthetic cathelicidin. Cultures were incubated for a further 72 hours.

### *In vitro* re-stimulation experiments

7 days post-immunisation, inguinal lymph nodes were isolated and prepared by mashing the tissue through a 100uM strainer and washing with PBS. 100,000 cells were plated per well in round-bottom 96-well plates in complete medium (RPMI, 10% foetal calf serum, 10 units/ml penicillin, 10 μg/ml streptomycin and 2 mM L-glutamine, all supplied by Gibco, ThermoFisher UK). All cells were incubated for 72 hours with Myelin Oligodendrocyte Protein (MOG) (2.5ug/ml; Sigma-Aldrich, M4939), with or without rIL-12 (25ng/ml) and cathelicidin (2.5μM).

### ELISA

Concentrations of mouse IL-23 (R&D Systems, #DY1887) and IL-12 (R&D Systems #M1270) were determined in spinal cord supernatants 7- and 14-days post immunisation by ELISA, as per the manufacturer’s guidelines.

### Immunohistochemistry mouse tissue

Mice were perfused with 4% paraformaldehyde (PFA) (Sigma-Aldrich, #158127) at indicated time points after immunisation. Whole brain, cervical and lumbar spinal cord, and inguinalr lymph nodes were isolated and fixed in 4% PFA overnight at 4°C. Brains and spinal cords were transferred to 15% sucrose for 24 hours, then 30% sucrose for 24 hours and embedded in Optimal Cutting Temperature (OCT) compound to be cryosectioned (5μm). Lymph nodes were transferred to 70% ethanol and embedded in paraffin.

Frozen sections were air dried for 10 minutes and paraffin-embedded slides were de-waxed and rehydrated. Haematoxylin and eosin (H&E) staining was carried out on tissue sections to understand inflammatory infiltration and the location of lesions within the spinal cord. For spinal cord sections, antigens were retrieved either by microwaving in tri-sodium citrate buffer, pH6 for 5 minutes or by 10 minutes incubation with 5mg/ml proteinase K (ThermoFisher UK, #AM2548) at 37 degrees. Slides were blocked with 25% donkey serum for 1h at RT, then primary antibodies were incubated in 10% donkey serum overnight at 4°C. Slides were then incubated with a combination of the following secondary antibodies: donkey anti-rat 488 (Invitrogen, #A21208), donkey anti-rabbit 555 (Invitrogen, #A31572), chicken anti-rat 647 (Invitrogen, #A31572) or chicken anti-goat 647 (Invitrogen, #214690) all at 1:1000 for 1 hour at RT. Slides were counterstained with Hoechst at 1:1000 (Abcam, #H1399). For lymph node sections, antigens were retrieved using 5mg/ml proteinase K, as above. Slides were blocked with 3% H_2_O_2_, 25% goat serum and avidin/biotin and the first primary antibody was added overnight at 4°C. The antigen was visualised using DAB Substrate kit (Vector Labs, #SK-4105) and subsequently blocked with Bloxall Endogenous Blocking Solution (Vector Labs, #SP- 6000-100) and 25% horse serum. The second primary antibody was added overnight at 4°C and visualised the following day using Vector Red Substrate Kit (Vector Labs, SK-5100). Slides were counterstained with haematoxylin. All stained sections were mounted in Fluoromount G, scanned on a ZEISS AxioScan.Z1 slide scanner and analysed using ZEISS ZEN software.

For quantification of markers in the spinal cord, 4 animals were analysed per time point. 5-10 sections per mouse were analysed across the whole spinal cord cross section, each datapoint on the graphs represent a cross-section analysed. For inguinal lymph nodes, 3 animals were analysed per time point. 2 whole lymph node cross sections per mouse were analysed, each datapoint on the graphs represent a cross-section analysed.

### Antibodies (mouse)

CRAMP (rabbit polyclonal, Innovagen, #PA-CRPL, 1:500-1:1000), Ly6G (IA8, Biolegend, #127601, 1:50), CD3 (17A2, Biolegend, #100209, 1:50), CD31 (rat polyclonal, R&D Systems, #AF3628, 1:100), peripheral node addressin (MECA-79, Novus Biologicals, #NB100- 77673SS, 1:100) and Iba1 (goat polyclonal, Abcam #ab5076, 1:500).

### Human post-mortem brain tissue

Post-mortem tissue from secondary progressive MS patients and control patients who died of non-neurological causes, with ethical approval and informed consent, were obtained from the UK Multiple Sclerosis Tissue Bank in collaboration with Professor Anna Williams, University of Edinburgh. Tissue was snap frozen and lesions classified as active, chronic active, chronic inactive and remyelinating using Luxol Fast Blue-Cresyl Violet staining and Oil Red O staining, according to the International Classification of Neurological Diseases.

### Immunohistochemistry human tissue

Frozen tissue sections were fixed in 4% paraformaldehyde and subsequently delipidised in methanol. Antigen retrieval was performed with heating in acid citric buffer. Sections were incubated overnight at 4°C with primary antibodies for CD31 or CD68 (antibodies listed below). On day 2 secondary antibodies were added for 30 minutes at room temperature and visualised with the Vector Blue Alkaline Phosphatase Substrate Kit (Vector Laboratories #SK-5300). Following this, sections were incubated overnight at 4°C with the second primary antibody for cathelicidin. On day 3 secondary antibodies were added and visualised with the Vector Red Alkaline Phosphatase Substrate Kit (Vector Laboratories #SK-5100). For neutrophil elastase (NE) and cathelicidin staining, frozen tissue sections were fixed in acetone for 5 minutes and delipidised in ethanol for 10 minutes. Antigens were retrieved by heat-induced sodium citrate and permeabilised with Triton X100. True black in 70% ethanol (Cambridge Bioscience #BT2300, dilution 1:20) was added to the slides for 20 seconds and washed thoroughly with PBS and subsequently blocked with 25% donkey serum. Primary antibodies were incubated overnight at 4°C. In all samples, nuclei were counterstained with Hoechst as above and slides mounted with Fluoromount G medium. Relevant IgG isotype antibodies and secondary antibodies alone were used as negative controls.

Cathelicidin-positive cells and NE cathelicidin-double positive cells were quantified in MS brains within each lesion type across the whole lesion and calculated as positive cells per cm^2^. The mean area of all lesions was calculated, and three white matter areas of this size were counted in control brains. The number of endothelial cells expressing cathelicidin was quantified as the % of CD31+ blood vessels expressing cathelicidin within a lesion.

### Antibodies (human)

Cathelicidin (rabbit polyclonal, abcam #ab69192, dilution 1:100), cathelicidin (rabbit polyclonal, abcam #ab69484, dilution 1:100), neutrophil elastase (clone NP57, Dako M0752, dilution 1:100), CD68 (KP1, Dako M0814, dilution 1:100) and CD31 (JC70A, Dako M0823, dilution 1:100).

## Data availability

All data are available from the corresponding author. The Nanostring dataset has been deposited in the GEO database with accession code GSE188655.

### Statistics

Two groups were compared with two-way un-paired or paired t test. Multiple groups were compared by one- or two-way ANOVA analysis. All data was analysed with Graphpad Prism. Sample sizes and what each data point on the graphs represent is detailed in the figure legends.

---

**Supplementary Figure 1:**
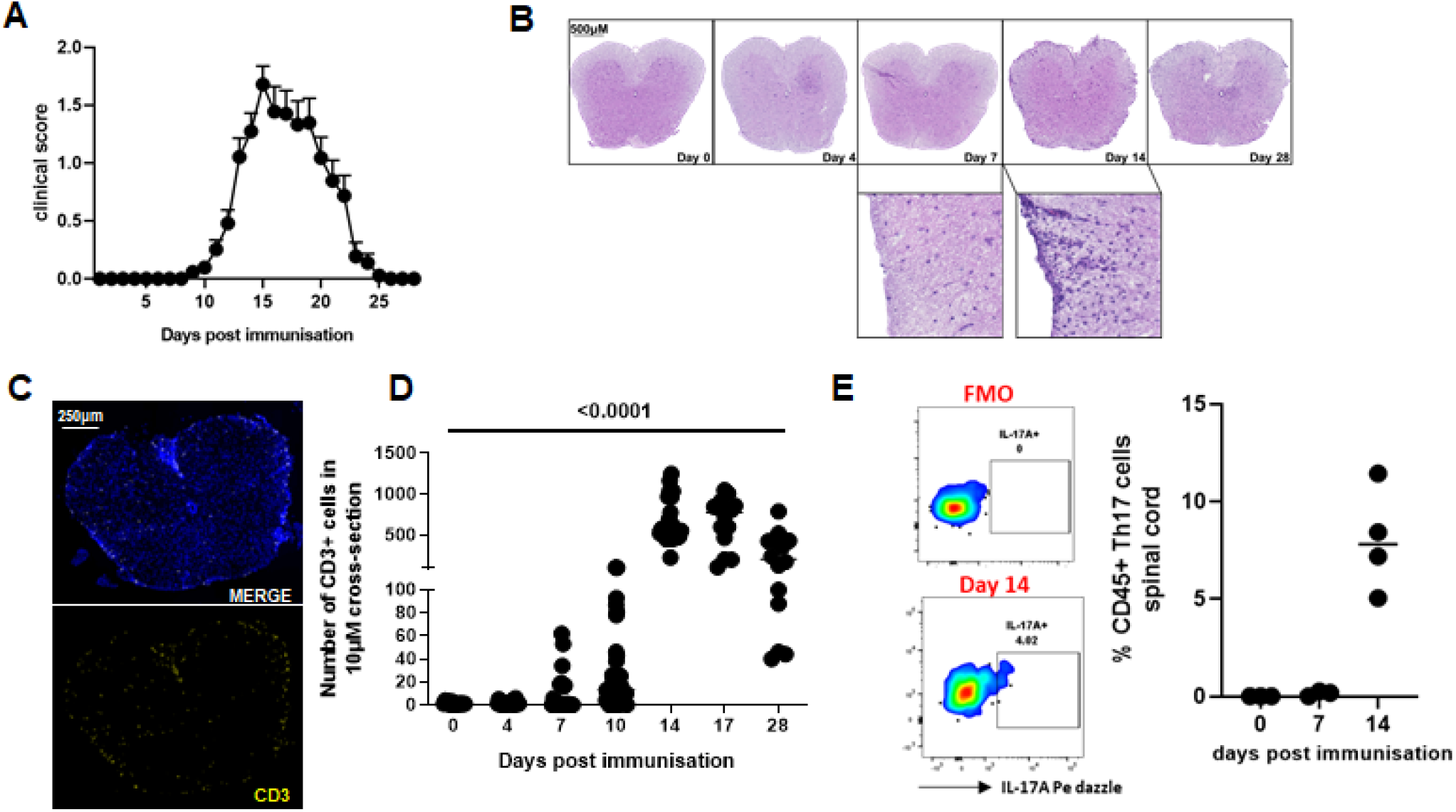
Establishment of the EAE model in wildtype mice. WT mice were immunised with myelin oligodendrocyte glycoprotein in complete Freund’s adjuvant and pertussis toxin was given. Mice were (A) tracked for clinical signs of disease over a time course and (B) at various times were perfused with 4% paraformaldehyde and spinal cords removed. Haematoxylin and eosin staining was performed to detect inflammatory infiltrate and (C, D) anti-CD3 immunofluorescent staining performed to detect T cell infiltrate. (E) Th17 cell numbers in the spinal cord were quantified by flow cytometry. Data shown are (A) mean with standard error and (D, E) individual data points with line at median. N values: A 85 mice; D 12-60 sections from 3-4 mice; E −4. Images are representative of 3 mice.

**Supplementary table 1:** Information relating to human tissue donors. MS− Multiple Sclerosis; SP − secondary progressive; PP − primary progressive.

| MS Patients | Sex | Age (Years) | MS Classification | Disease Duration (Years) | Post-mortem Interval (Hours) |
| --- | --- | --- | --- | --- | --- |
| MS100 | M | 46 | SP | 8 | 7 |
| MS136 | M | 40 | SP | 9 | 10 |
| MS154 | F | 34 | SP | 21 | 12 |
| MS176 | M | 37 | PP | 27 | 12 |
| MS207 | F | 46 | SP | 25 | 10 |
| MS230 | F | 42 | SP | 19 | 31 |
| MS122 | M | 44 | SP | Not known | 16 |

| Controls | Sex | Age (Years) | Cause of Death | Post-mortem Interval |
| --- | --- | --- | --- | --- |
| CO25 | M | 35 | Carcinoma of the tongue | 22 |
| CO28 | F | 60 | Ovarian Cancer | 13 |
| CO39 | M | 82 | Myelodysplastic syndrome, Rheumatoid arthritis | 21 |

**Supplementary Figure 2.**
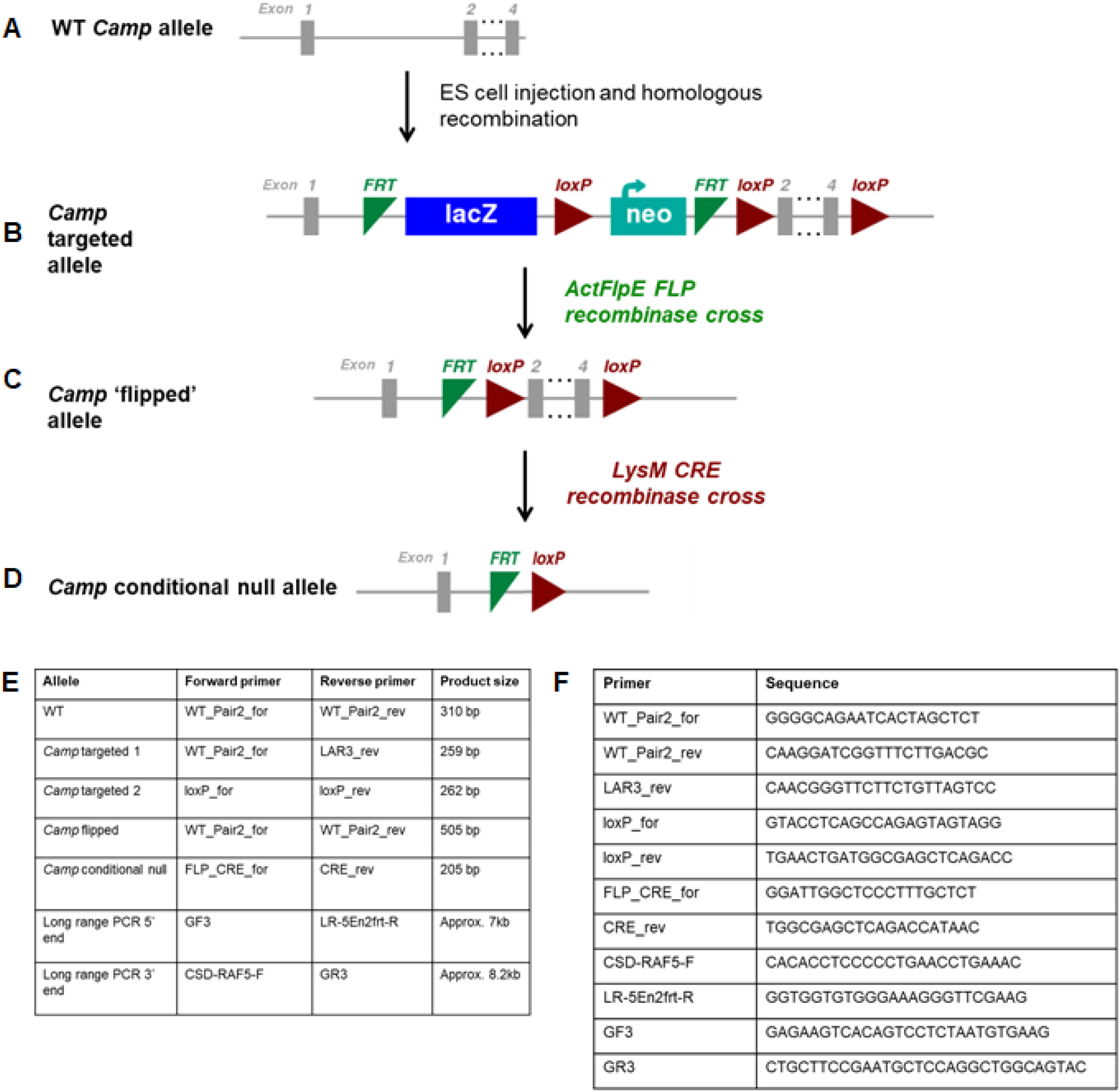
Generation and validation of the LysMere mouse. The WT *Camp* gene (A) consists of 4 exons on chromosome 9. *Camp*^*tm1a(EUCOMM) Hmgo*^ ES cells (JM8A3.N1; cell clone ID HEPD0722_1_E10; MGl:4950203) cells targeting the *Camp* locus were purchased from EUeOMM, injected into e57/BI6J blastocyst stage embryos and subsequently transferred to recipient female mice. Male chimeric progeny were mated with e57B1/6JerI female mice to establish germ line transmission, confirmed by short range and long range PeR (Primer combinations and sequences in Methods). Targeted mice *(Camp*^*tm1a(EUCOMM) Hmgo;*^ B) were then crossed with ActFlpE (SJL Tg(AeTFLPe)9205Dym/J) mice on e57BI6/JerI bacKground mice to generate mice with a ‘flipped’ allele *(Camp*^*tm1a(EUCOMM) Hmgo / ACTFLPe*^; C) lacKing the lacZ and neo vector cassettes, intercrossed to hOmozygosity and confirmed by PeR. Confirmation of ere recombination to generate a conditional null allele (with excision of *Camp* exons 2 − 4) was provided by administration **of** soluble tat ere recombinase to one cell *Camp*^*tm1a(EUCOMM) Hmgo / ACTFLPe*^ embryos obtained by IVF, transferred into recipients at 2.5 days, followed by PeR confirmation and sequencing of the resulting conditional null allele on E12 embryos. *Camp*^*tm1a(EucoMM) Hmgo / ACTFLPe*^ were crossed with a myeloid-specific CRE recombinase line LysMere(Tg (Lyz2tm1(cre)lfo) to generate the *Camp* conditional null mice *(Camp*^*tm1a(EUCOMM) Hmgo I ACTFLPe I Tg (Lyz2fm1(cre)lfo;*^ D) designated *LysMCreFLCamp* mice and bred to congenicity for n=1o generations. Breeding pairs hOmozygous for floxed *Camp* were heterozygous or wild type for LysMere (designated *LysMCreFLCamp* and *FLCamp* respectively) and studied experimentally compared to littermates controls.. (E) primers and (F) sequences used.

## References

1. Dendrou CA, Fugger L, Friese MA. Immunopathology of multiple sclerosis. Nat Rev Immunol. 2015;15(9):545–58.

2. Ivanov II, McKenzie BS, Zhou L, Tadokoro CE, Lepelley A, Lafaille JJ, et al. The Orphan Nuclear Receptor RORγt Directs the Differentiation Program of Proinflammatory IL-17+ T Helper Cells. Cell. 2006;126(6):1121–33.

3. Kurschus FC. T cell mediated pathogenesis in EAE: Molecular mechanisms. Biomed J. 2015;38(3):183–93.

4. Carlson T, Kroenke M, Rao P, Lane TE, Segal B. The Th17-ELR+ CXC chemokine pathway is essential for the development of central nervous system autoimmune disease. J Exp Med. 2008;205(4):811–23.

5. Tzartos JS, Friese MA, Craner MJ, Palace J, Newcombe J, Esiri MM, et al. Interleukin-17 production in central nervous system-infiltrating T cells and glial cells is associated with active disease in multiple sclerosis. Am J Pathol. 2008;172(1):146–55.

6. Stromnes IM, Goverman JM. Passive induction of experimental allergic encephalomyelitis. Nat Protoc. 2006;1(4):1952–60.

7. Ben-Nun A, Wekerle H. and J. R. Cohen. 1981. The rapid isolation of clonable antigenspecific T lymphocyte lines capable of mediating autoimmune encephalomyelitis. Eur J Immunol. 11:195.

8. 1. McPherson RC, Cambrook HE, O’Connor RA, Anderton SM. Induction of passive EAE using myelin-reactive CD4+ T cells. Methods Mol Biol. 2014;1193:187–98.

9. Yasuda K, Takeuchi Y, Hirota K. The pathogenicity of Th17 cells in autoimmune diseases. Semin Immunopathol. 2019;41(3):283–97.

10. Ifergan I, Kebir H, Alvarez JI, Marceau G, Bernard M, Bourbonnière L, et al. Central nervous system recruitment of effector memory CD8+ T lymphocytes during neuroinflammation is dependent on α4 integrin. Brain. 2011;134(12):3560–77.

11. Hirota K, Duarte JH, Veldhoen M, Hornsby E, Li Y, Cua DJ, et al. Fate mapping of IL-17-producing T cells in inflammatory responses. Nat Immunol. 2011;12(3):255–63.

12. Komiyama Y, Nakae S, Matsuki T, Nambu A, Ishigame H, Kakuta S, et al. IL-17 plays an important role in the development of experimental autoimmune encephalomyelitis. J Immunol. 2006;177(1):566–73.

13. McGinley AM, Sutton CE, Edwards SC, Leane CM, DeCourcey J, Teijeiro A, et al. Interleukin-17A Serves a Priming Role in Autoimmunity by Recruiting IL-1β-Producing Myeloid Cells that Promote Pathogenic T Cells. Immunity. 2020;52(2):342–56.e6.

14. Olsson T. Cytokines in neuroinflammatory disease: role of myelin autoreactive T cell production of interferon-gamma. Journal of neuroimmunology. 1992;40(2-3):211–8.

15. Ley K. The second touch hypothesis: T cell activation, homing and polarization. F1000Res. 2014;3:37.

16. Furtado GC, Marcondes MCG, Latkowski J-A, Tsai J, Wensky A, Lafaille JJ. Swift entry of myelin-specific T lymphocytes into the central nervous system in spontaneous autoimmune encephalomyelitis. The Journal of Immunology. 2008;181(7):4648–55.

17. Stoll S, Delon J, Brotz TM, Germain RN. Dynamic imaging of T cell-dendritic cell interactions in lymph nodes. Science. 2002;296(5574):1873–6.

18. Langrish CL, Chen Y, Blumenschein WM, Mattson J, Basham B, Sedgwick JD, et al. IL-23 drives a pathogenic T cell population that induces autoimmune inflammation. J Exp Med. 2005;201(2):233–40.

19. Simmons SB, Liggitt D, Goverman JM. Cytokine-regulated neutrophil recruitment is required for brain but not spinal cord inflammation during experimental autoimmune encephalomyelitis. J Immunol. 2014;193(2):555–63.

20. Pierson ER, Wagner CA, Goverman JM. The contribution of neutrophils to CNS autoimmunity. Clin Immunol. 2018;189:23–8.

21. Aubé B, Lévesque SA, Paré A, Chamma É, Kébir H, Gorina R, et al. Neutrophils mediate blood-spinal cord barrier disruption in demyelinating neuroinflammatory diseases. J Immunol. 2014;193(5):2438–54.

22. Song J, Wu C, Korpos E, Zhang X, Agrawal SM, Wang Y, et al. Focal MMP-2 and MMP-9 activity at the blood-brain barrier promotes chemokine-induced leukocyte migration. Cell Rep. 2015;10(7):1040–54.

23. Ouyang W, Kolls JK, Zheng Y. The biological functions of T helper 17 cell effector cytokines in inflammation. Immunity. 2008;28(4):454–67.

24. Pelletier M, Maggi L, Micheletti A, Lazzeri E, Tamassia N, Costantini C, et al. Evidence for a cross-talk between human neutrophils and Th17 cells. Blood, The Journal of the American Society of Hematology. 2010;115(2):335–43.

25. Grist JJ, Marro BS, Skinner DD, Syage AR, Worne C, Doty DJ, et al. Induced CNS expression of CXCL1 augments neurologic disease in a murine model of multiple sclerosis via enhanced neutrophil recruitment. Eur J Immunol. 2018;48(7):1199–210.

26. Haschka D, Tymoszuk P, Bsteh G, Petzer V, Berek K, Theurl I, et al. Expansion of Neutrophils and Classical and Nonclassical Monocytes as a Hallmark in Relapsing-Remitting Multiple Sclerosis. Front Immunol. 2020;11:594.

27. Woodberry T, Bouffler SE, Wilson AS, Buckland RL, Brüstle A. The Emerging Role of Neutrophil Granulocytes in Multiple Sclerosis. J Clin Med. 2018;7(12).

28. 1. Naegele M, Tillack K, Reinhardt S, Schippling S, Martin R, Sospedra M. Neutrophils in multiple sclerosis are characterized by a primed phenotype. J Neuroimmunol. 2012;242(1-2):60–71.

29. Rumble JM, Huber AK, Krishnamoorthy G, Srinivasan A, Giles DA, Zhang X, et al. Neutrophil-related factors as biomarkers in EAE and MS. J Exp Med. 2015;212(1):23–35.

30. Steinbach K, Piedavent M, Bauer S, Neumann JT, Friese MA. Neutrophils amplify autoimmune central nervous system infiltrates by maturing local APCs. J Immunol. 2013;191(9):4531–9.

31. Minns D, Smith KJ, Alessandrini V, Hardisty G, Melrose L, Jackson-Jones L, et al. The neutrophil antimicrobial peptide cathelicidin promotes Th17 differentiation. Nature Communications. 2021;12(1):1285.

32. Zorzella-Pezavento SFG, Chiuso-Minicucci F, França TGD, Ishikawa LLW, da Rosa LC, Marques C, et al. Persistent Inflammation in the CNS during Chronic EAE Despite Local Absence of IL-17 Production. Mediators of Inflammation. 2013;2013:519627.

33. Wu F, Cao W, Yang Y, Liu A. Extensive infiltration of neutrophils in the acute phase of experimental autoimmune encephalomyelitis in C57BL/6 mice. Histochem Cell Biol. 2010;133(3):313–22.

34. Gudmundsson GH, Agerberth B, Odeberg J, Bergman T, Olsson B, Salcedo R. The human gene FALL39 and processing of the cathelin precursor to the antibacterial peptide LL-37 in granulocytes. Eur J Biochem. 1996;238(2):325–32.

35. Sørensen OE, Follin P, Johnsen AH, Calafat J, Tjabringa GS, Hiemstra PS, et al. Human cathelicidin, hCAP-18, is processed to the antimicrobial peptide LL-37 by extracellular cleavage with proteinase 3. Blood. 2001;97(12):3951–9.

36. Lande R, Ganguly D, Facchinetti V, Frasca L, Conrad C, Gregorio J, et al. Neutrophils activate plasmacytoid dendritic cells by releasing self-DNA-peptide complexes in systemic lupus erythematosus. Sci Transl Med. 2011;3(73):73ra19.

37. Dudal S, Turriere C, Bessoles S, Fontes P, Sanchez F, Liautard J, et al. Release of LL-37 by Activated Human Vγ9Vδ2 T Cells: A Microbicidal Weapon against *Brucella suis*. The Journal of Immunology. 2006;177(8):5533.

38. Agerberth B, Charo J, Werr J, Olsson B, Idali F, Lindbom L, et al. The human antimicrobial and chemotactic peptides LL-37 and alpha-defensins are expressed by specific lymphocyte and monocyte populations. Blood. 2000;96(9):3086–93.

39. Zhang LJ, Guerrero-Juarez CF, Hata T, Bapat SP, Ramos R, Plikus MV, et al. Innate immunity. Dermal adipocytes protect against invasive Staphylococcus aureus skin infection. Science. 2015;347(6217):67–71.

40. Yim S, Dhawan P, Ragunath C, Christakos S, Diamond G. Induction of cathelicidin in normal and CF bronchial epithelial cells by 1,25-dihydroxyvitamin D(3). J Cyst Fibros. 2007;6(6):403–10.

41. Bals R, Wang X, Zasloff M, Wilson JM. The peptide antibiotic LL-37/hCAP-18 is expressed in epithelia of the human lung where it has broad antimicrobial activity at the airway surface. Proceedings of the National Academy of Sciences. 1998;95(16):9541.

42. Frohm Nilsson M, Sandstedt B, Sørensen O, Weber G, Borregaard N, Ståhle-Bäckdahl M. The human cationic antimicrobial protein (hCAP18), a peptide antibiotic, is widely expressed in human squamous epithelia and colocalizes with interleukin-6. Infect Immun. 1999;67(5):2561–6.

43. Di Nardo A, Braff MH, Taylor KR, Na C, Granstein RD, McInturff JE, et al. Cathelicidin Antimicrobial Peptides Block Dendritic Cell TLR4 Activation and Allergic Contact Sensitization. The Journal of Immunology. 2007;178(3):1829.

44. Elkjaer ML, Frisch T, Reynolds R, Kacprowski T, Burton M, Kruse TA, et al. Molecular signature of different lesion types in the brain white matter of patients with progressive multiple sclerosis. Acta Neuropathologica Communications. 2019;7(1):205.

45. Frischer JM, Weigand SD, Guo Y, Kale N, Parisi JE, Pirko I, et al. Clinical and pathological insights into the dynamic nature of the white matter multiple sclerosis plaque. Ann Neurol. 2015;78(5):710–21.

46. Reynolds R, Roncaroli F, Nicholas R, Radotra B, Gveric D, Howell O. The neuropathological basis of clinical progression in multiple sclerosis. Acta Neuropathol. 2011;122(2):155–70.

47. Nizet V, Ohtake T, Lauth X, Trowbridge J, Rudisill J, Dorschner RA, et al. Innate antimicrobial peptide protects the skin from invasive bacterial infection. Nature. 2001;414(6862):454–7.

48. De Y, Chen Q, Schmidt AP, Anderson GM, Wang JM, Wooters J, et al. LL-37, the neutrophil granule-and epithelial cell-derived cathelicidin, utilizes formyl peptide receptor-like 1 (FPRL1) as a receptor to chemoattract human peripheral blood neutrophils, monocytes, and T cells. The Journal of experimental medicine. 2000;192(7):1069–74.

49. Ghoreschi K, Laurence A, Yang X-P, Tato CM, McGeachy MJ, Konkel JE, et al. Generation of pathogenic TH17 cells in the absence of TGF-β signalling. Nature. 2010;467(7318):967–71.

50. Zamvil S, Nelson P, Trotter J, Mitchell D, Knobler R, Fritz R, et al. T-cell clones specific for myelin basic protein induce chronic relapsing paralysis and demyelination. Nature. 1985;317(6035):355–8.

51. Goverman J. Autoimmune T cell responses in the central nervous system. Nat Rev Immunol. 2009;9(6):393–407.

52. Reboldi A, Coisne C, Baumjohann D, Benvenuto F, Bottinelli D, Lira S, et al. C-C chemokine receptor 6-regulated entry of TH-17 cells into the CNS through the choroid plexus is required for the initiation of EAE. Nat Immunol. 2009;10(5):514–23.

53. Baron JL, Madri JA, Ruddle NH, Hashim G, Janeway CA, Jr. Surface expression of alpha 4 integrin by CD4 T cells is required for their entry into brain parenchyma. J Exp Med. 1993;177(1):57–68.

54. Leonard J, Waldburger K, Goldman S. Prevention of experimental autoimmune encephalomyelitis by antibodies against interleukin 12. Journal of Experimental Medicine. 1995;181(1):381–6.

55. Segal BM, Shevach EM. IL-12 unmasks latent autoimmune disease in resistant mice. The Journal of experimental medicine. 1996;184(2):771–5.

56. Yang XO, Chang SH, Park H, Nurieva R, Shah B, Acero L, et al. Regulation of inflammatory responses by IL-17F. Journal of Experimental Medicine. 2008;205(5):1063–75.

57. Lee YK, Turner H, Maynard CL, Oliver JR, Chen D, Elson CO, et al. Late developmental plasticity in the T helper 17 lineage. Immunity. 2009;30(1):92–107.

58. Lee YK, Mukasa R, Hatton RD, Weaver CT. Developmental plasticity of Th17 and Treg cells. Curr Opin Immunol. 2009;21(3):274–80.

59. Yang Y, Weiner J, Liu Y, Smith AJ, Huss DJ, Winger R, et al. T-bet is essential for encephalitogenicity of both Th1 and Th17 cells. J Exp Med. 2009;206(7):1549–64.

60. Kurschus FC, Croxford AL, Heinen AP, Wörtge S, Ielo D, Waisman A. Genetic proof for the transient nature of the Th17 phenotype. Eur J Immunol. 2010;40(12):3336–46.

61. Kebir H, Ifergan I, Alvarez JI, Bernard M, Poirier J, Arbour N, et al. Preferential recruitment of interferon-γ–expressing TH17 cells in multiple sclerosis. Annals of Neurology: Official Journal of the American Neurological Association and the Child Neurology Society. 2009;66(3):390–402.

62. O’Connor RA, Prendergast CT, Sabatos CA, Lau CWZ, Leech MD, Wraith DC, et al. Cutting edge: Th1 cells facilitate the entry of Th17 cells to the central nervous system during experimental autoimmune encephalomyelitis. Journal of immunology (Baltimore, Md : 1950). 2008;181(6):3750–4.

63. Harbour SN, Maynard CL, Zindl CL, Schoeb TR, Weaver CT. Th17 cells give rise to Th1 cells that are required for the pathogenesis of colitis. Proceedings of the National Academy of Sciences. 2015;112(22):7061–6.

64. Duhen R, Glatigny S, Arbelaez CA, Blair TC, Oukka M, Bettelli E. Cutting edge: The pathogenicity of IFN-γ–producing Th17 cells is independent of T-bet. The Journal of Immunology. 2013;190(9):4478–82.

65. Duhen T, Campbell DJ. IL-1β promotes the differentiation of polyfunctional human CCR6+CXCR3+ Th1/17 cells that are specific for pathogenic and commensal microbes. J Immunol. 2014;193(1):120–9.

66. Takahashi T, Kulkarni NN, Lee EY, Zhang LJ, Wong GCL, Gallo RL. Cathelicidin promotes inflammation by enabling binding of self-RNA to cell surface scavenger receptors. Sci Rep. 2018;8(1):4032.

67. Hiemstra PS, Amatngalim GD, van der Does AM, Taube C. Antimicrobial Peptides and Innate Lung Defenses: Role in Infectious and Noninfectious Lung Diseases and Therapeutic Applications. Chest. 2016;149(2):545–51.

68. Lande R, Gregorio J, Facchinetti V, Chatterjee B, Wang Y-H, Homey B, et al. Plasmacytoid dendritic cells sense self-DNA coupled with antimicrobial peptide. Nature. 2007;449(7162):564–9.

69. Takahashi T, Gallo RL. The Critical and Multifunctional Roles of Antimicrobial Peptides in Dermatology. Dermatol Clin. 2017;35(1):39–50.

70. Kahlenberg JM, Kaplan MJ. Little Peptide, Big Effects: The Role of LL-37 in Inflammation and Autoimmune Disease. The Journal of Immunology. 2013;191(10):4895–901.

71. McHugh BJ, Wang R, Li HN, Beaumont PE, Kells R, Stevens H, et al. Cathelicidin is a “fire alarm”, generating protective NLRP3-dependent airway epithelial cell inflammatory responses during infection with Pseudomonas aeruginosa. PLoS Pathog. 2019;15(4):e1007694.

72. Bergman P, Termén S, Johansson L, Nyström L, Arenas E, Jonsson AB, et al. The antimicrobial peptide rCRAMP is present in the central nervous system of the rat. J Neurochem. 2005;93(5):1132–40.

73. Wan M, Sabirsh A, Wetterholm A, Agerberth B, Haeggström JZ. Leukotriene B4 triggers release of the cathelicidin LL-37 from human neutrophils: novel lipid-peptide interactions in innate immune responses. FASEB J. 2007;21(11):2897–905.

74. Kulkarni NN, Yi Z, Huehnken C, Agerberth B, Gudmundsson GH. Phenylbutyrate induces cathelicidin expression via the vitamin D receptor: Linkage to inflammatory and growth factor cytokines pathways. Mol Immunol. 2015;63(2):530–9.

75. Bergman P, Johansson L, Wan H, Jones A, Gallo RL, Gudmundsson GH, et al. Induction of the antimicrobial peptide CRAMP in the blood-brain barrier and meninges after meningococcal infection. Infect Immun. 2006;74(12):6982–91.

76. Xu X, Cai X, Zhu Y, He W, Wu Q, Shi X, et al. MFG-E8 inhibits Aβ-induced microglial production of cathelicidin-related antimicrobial peptide: A suitable target against Alzheimer’s disease. Cellular Immunology. 2018;331:59–66.

77. Brandenburg LO, Varoga D, Nicolaeva N, Leib SL, Wilms H, Podschun R, et al. Role of glial cells in the functional expression of LL-37/rat cathelin-related antimicrobial peptide in meningitis. J Neuropathol Exp Neurol. 2008;67(11):1041–54.

78. Lee M, Shi X, Barron AE, McGeer E, McGeer PL. Human antimicrobial peptide LL-37 induces glial-mediated neuroinflammation. Biochemical Pharmacology. 2015;94(2):130–41.

